# On measuring selection in cancer from subclonal mutation frequencies

**DOI:** 10.1101/529396

**Authors:** Ivana Bozic, Chay Paterson, Bartlomiej Waclaw

## Abstract

Recently available cancer sequencing data have revealed a complex view of the cancer genome containing a multitude of mutations, including drivers responsible for cancer progression and neutral passengers. Measuring selection in cancer and distinguishing drivers from passengers have important implications for development of novel treatment strategies. It has recently been argued that a third of cancers are evolving neutrally, as their mutational frequency spectrum follows a 1/*f* power law expected from neutral evolution in a particular intermediate frequency range. We study a stochastic model of cancer evolution and derive a formula for the probability distribution of the cancer cell frequency of a subclonal driver, demonstrating that driver frequency is biased towards 0 and 1. We show that it is difficult to capture a driver mutation at an intermediate frequency, and thus the calling of neutrality due to a lack of such driver will significantly overestimate the number of neutrally evolving tumors. Our approach provides precise quantification of the validity of the 1/*f* statistic across the entire range of all relevant parameter values. Our results are also applicable to the question of distinguishing driver and passenger mutations in a general exponentially expanding population.

## INTRODUCTION

Darwinian evolution in cancer has been the subject of intense research in the past decade. In particular, the problem of distinguishing driver mutations that carry a selective advantage from passenger mutations, and their role in shaping intra-tumor genetic heterogeneity has come to the fore^1-5^. Determining which mutations in cancer are drivers and which are passengers is one of the most pressing questions in cancer genomics, as identification of new driver mutations can contribute to development of new targeted therapeutics^6,7^ and personalized medicine^8^. Numerous methods for classifying driver and passenger mutations and measuring selection in cancer have been developed, including those that identify driver genes based on how frequently they are mutated^2^, specific mutation patterns^9,10^, and dN/dS ratios^1,11^. These methods can reliably identify driver genes mutated in an high proportion of tumors of a given type (>20%); using such methods to find less common drivers would require a large number of cancer samples^12^, and drivers unique to a single or a small number of patients could still be missed.

Several recent papers attempt to measure the magnitude of selection operating during cancer evolution using the frequency distribution of subclonal mutations in an individual patient’s cancer. In a seminal paper, Williams et al. used mutant allele frequencies to conclude that a significant fraction (∼1/3) of cancers evolve neutrally^13^. Subsequent studies focused on quantifying the strength of selection and distinguishing it from “effectively neutral” cancer evolution^14,15^. These works are based upon the assumption that drivers that arose after cancer initiation will be present at a macroscopic but clearly subclonal frequency (i.e. “detectable”), which will make the cumulative mutant allele frequency look different to the 1/*f* power law expected from neutral evolution^13,15^. Here we use a branching process model of cancer evolution to derive a formula for the probability of detection of a subclonal driver, and test the validity of the proposed 1/*f* statistic across all relevant parameter combinations.

## RESULTS

We consider a two-type stochastic model of cancer evolution (Fig. 1a). In the model, cancer is initiated by a single transformed cell. Progeny of this cell follow a branching process with birth rate *b* and death rate *d*. We set *r* = *b* – *d* > 0, so the population grows if it survives initial stochastic fluctuations. In addition, cancer cells can obtain a new driver mutation with rate *u* (Materials and Methods). Cells with the driver mutation replicate with rate *b*_1_ and die with rate *d*_1_ smaller than *b*_1_ (Fig. 1a,b). The subpopulation of driver-carrying cells has therefore a net growth rate *r*_1_ = *b*_1_ – *d*_1_, and we assume that *r*_1_ > *r* so that the additional driver increases the net growth rate by the factor *c* = *r*_1_/*r* > 1. We define *g* = *c* - 1 as the relative increase in the growth rate due to the driver.

**Figure 1.**
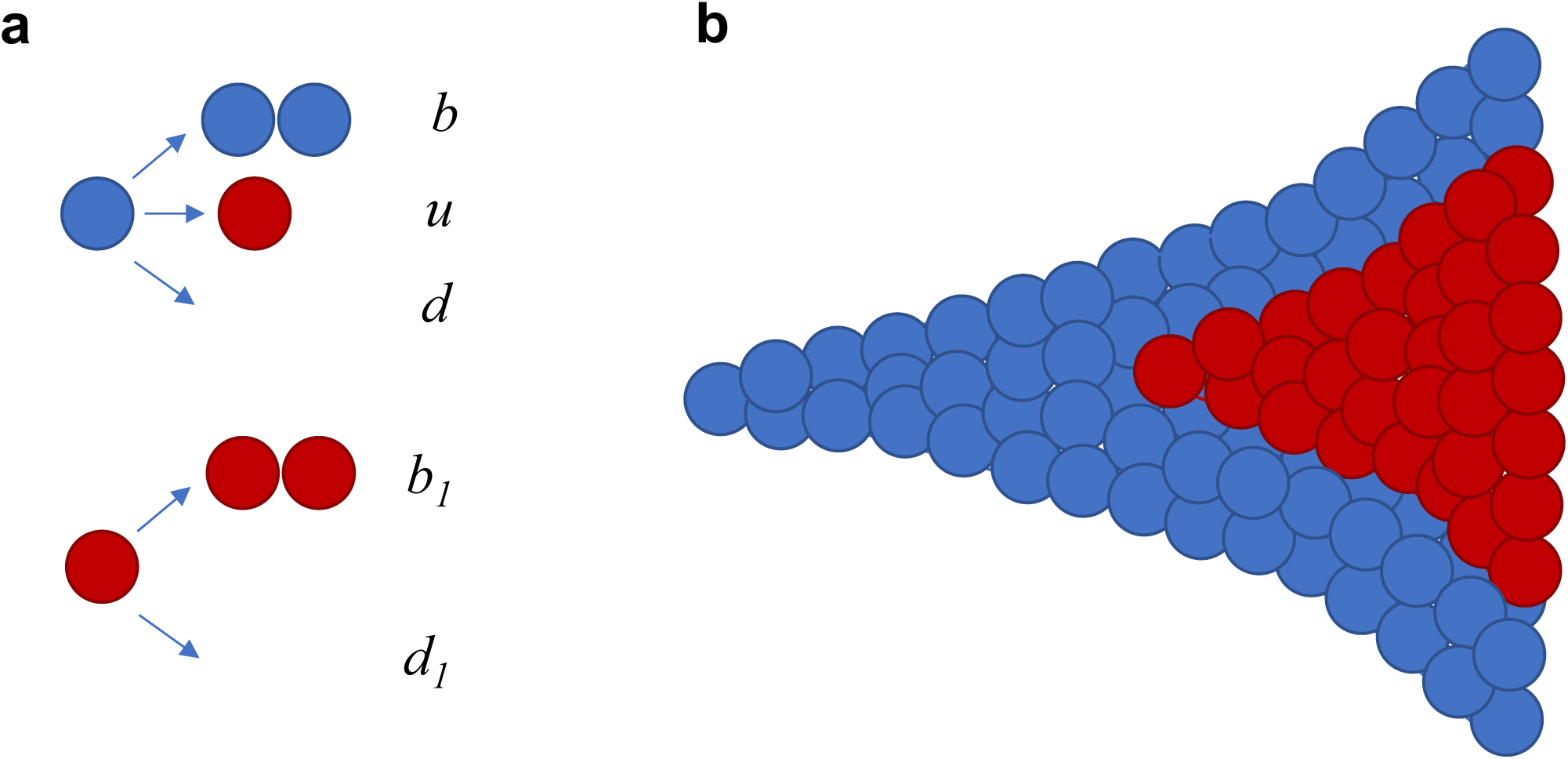
Schematic representation of the stochastic model of tumor evolution. **a,** Parental cells (blue) divide with rate *b*, obtain an additional driver with rate *u*, and die with rate *d*. Cells with the additional driver (red) divide with rate *b*_1_ and die with rate *d*_1_. The ratio of net growth rates of cells with and without the driver, *c*=(*b*_1_-*d*_1_)/(*b*-*d*) is greater than 1. **b,** Growth begins with a single parental cell. We are interested in the fraction of cells with the driver as a function of the total number of tumor cells *M*.

We are interested in the frequency of cancer cells that carry the driver mutation. In a neutral process (*g* = 0), mutation frequency stabilizes and remains approximately constant at large times^16^. For *g* > 0 (driver with a selective advantage), the frequency of cells with the driver increases from ≈0 to ≈100% during tumor expansion. We derive the formula for the cumulative frequency distribution of a subclonal driver (Materials and Methods)

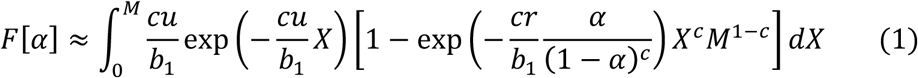

Here *F*[*α*]= *P*[*f* ≤ α] denotes the probability that subclonal driver frequency *f* is smaller than. *α. M* is number if cells in the tumor, *b*_1_ birth rate of cells with the driver, *r* net growth rate of cells without the driver, *u* driver mutation rate and *c*=1+*g* is the ratio of net growth rates of cells with and without the driver. Formula (1) is in excellent agreement with exact computer simulations of the branching process (Materials and Methods). We assume that a subclonal driver mutation can be detected, and able to skew the 1/*f* power law expected from neutral evolution, when its cancer cell frequency is between 20% and 80%. This range is much wider than the range 24% to 48% used in Williams et al.^13^ (mutant allele frequency range 12% to 24%). Thus, the probability that a driver can be detected is given by

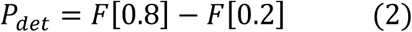

For moderate levels of selection, e.g., when the additional driver mutation increases the growth rate by *g* = 30%, the probability that the driver mutation is in the detectable range ([0.2,0.8]) is <15% for population sizes up to *M*=10^9^ cells, and remains below one third for *M* ≤ 10^11^ cells (Fig. 2a). For other cases considered here (70% and 100% increase in the net growth rate), the chance of detecting the subclonal driver is always <60% and - for a broad range of tumor sizes – less than 30%. The parameters used in Fig. 2a are from Bozic et al.^17^, and are typical for a moderately aggressive cancer (net growth rate 0.01/day). We show that the situation is qualitatively similar for faster and slower growing cancers in Figs. 2b,c. In summary, for moderate levels of selection (*g* = 30%), the chance of detecting a subclonal driver is small for almost any tumor size, and very strong selection (*g* = 100%) will be detectable only in small cancers. Strong selection (*g* = 70%) will be detectable at intermediate-size, moderately growing tumors; large, fast-growing tumors; or small, slow-growing tumors. Most notably, for all parameter values, even in the parameter regimes where the probability of detecting the subclonal driver is the highest, it is still below 60% (Fig. 2a-c).

**Figure 2.**
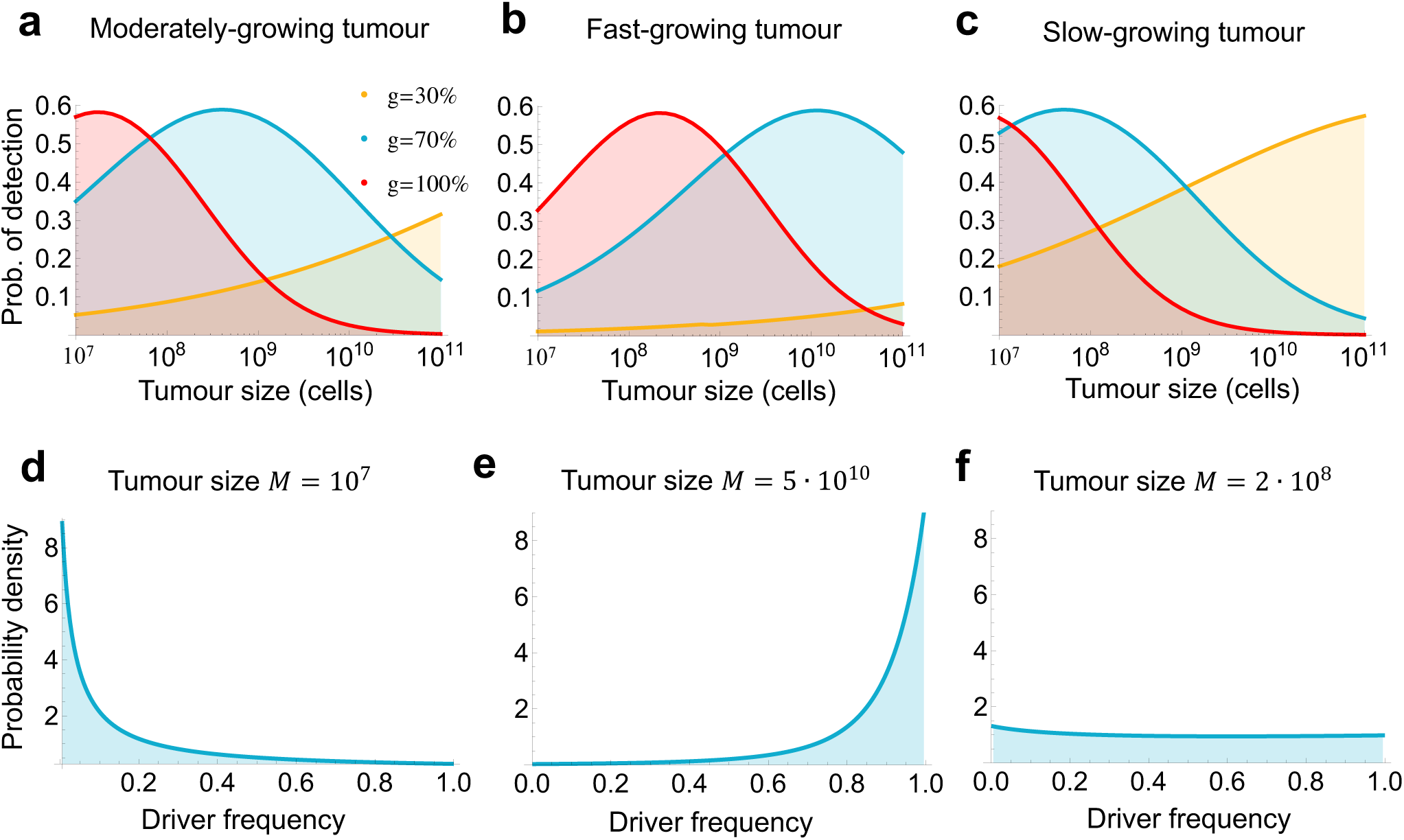
Frequency of a subclonal driver is biased towards 0 and 1. **a, b, c,** Probability that a subclonal driver is in the detectable range (0.2 ≤ *f*_*sub*_ ≤ 0.8) and thus able to skew the distribution of mutation frequencies expected from neutral evolution for three parameter regimes. For each parameter regime, we depict three levels of selection: moderate selection (driver increases net growth rate by *g* = 30%), strong selection (*g* = 70%), and very strong selection (*g* = 100%).Parameter values for **a,** moderately growing tumor^17^: *b*=0.14, *r=*0.01; **b,** fast growing tumor^23^: *b*=0.25, *r*=0.07; **c,** slow-growing tumor^18^: *b*=0.33, *r*=0.0013. Driver mutation rate^18^ *u*=10^-5^. All rates are per day and *b*=*b*_1_. **d, e, f,** Probability density for frequency of subclonal driver that increases the net growth rate by 70% in a moderately growing tumor (**a**). **d,** Driver frequency is biased towards 0 when tumor size is small. **e,** When tumor size is large, driver frequency is biased towards 1. **f,** When detection is most likely (at intermediate size), driver frequency distribution is almost flat.

Our model suggests that detecting a subclonal driver, and thus any deviation from neutral evolution, is generally difficult. The reason is that the frequency of cells with the new driver is biased toward 0 and 1. When the tumor is small, the fraction of driver-carrying cells is very close to zero, as there has not yet been enough time for the fitter subpopulation to expand (Fig. 2d). In contrast, for large tumors, driver-carrying subpopulation has already expanded and completely dominates the population, so its frequency is close to 100% (Fig. 2e). Interestingly, for sizes at which the chance of detecting the subclonal driver is highest (close to 60%), the frequency distribution is almost flat (Fig. 2f).

In Fig. 2 we used a previously estimated driver mutation rate *u*=10^-5^ per day^18^. To explore the effect of a higher or lower driver mutation rate on our conclusions, we first used recently published genomic data to determine an upper bound (10^-3^) and a lower bound (10^-7^ per day) on the driver mutation rate (Materials and Methods). We next performed a numerical grid search on the space of all parameters (driver mutation rate *u*, relative growth rate advantage of a driver C, net growth rate of tumor cells *r*, division rate of cells with the driver *b*_1_, and final number of tumor cells *M*). A wide range of values is taken for each parameter, including driver mutation rate *u* between 10^-7^ and 10^-3^ per day, and growth advantage of a subclonal driver C between 1% and 200% (see Materials and Methods for more details). The grid search demonstrated that the probability of detection of a subclonal driver is always below 60%, and that subclonal driver frequency is biased towards 0 and 1 across the entire range of reasonable parameter values of the carcinogenic process (Materials and Methods, Fig. 3). The intuitive reason behind this result is that the probability density function for subclonal driver frequency is convex across this entire parameter range (examples in Fig. 2d,e,f).

**Figure 3.**
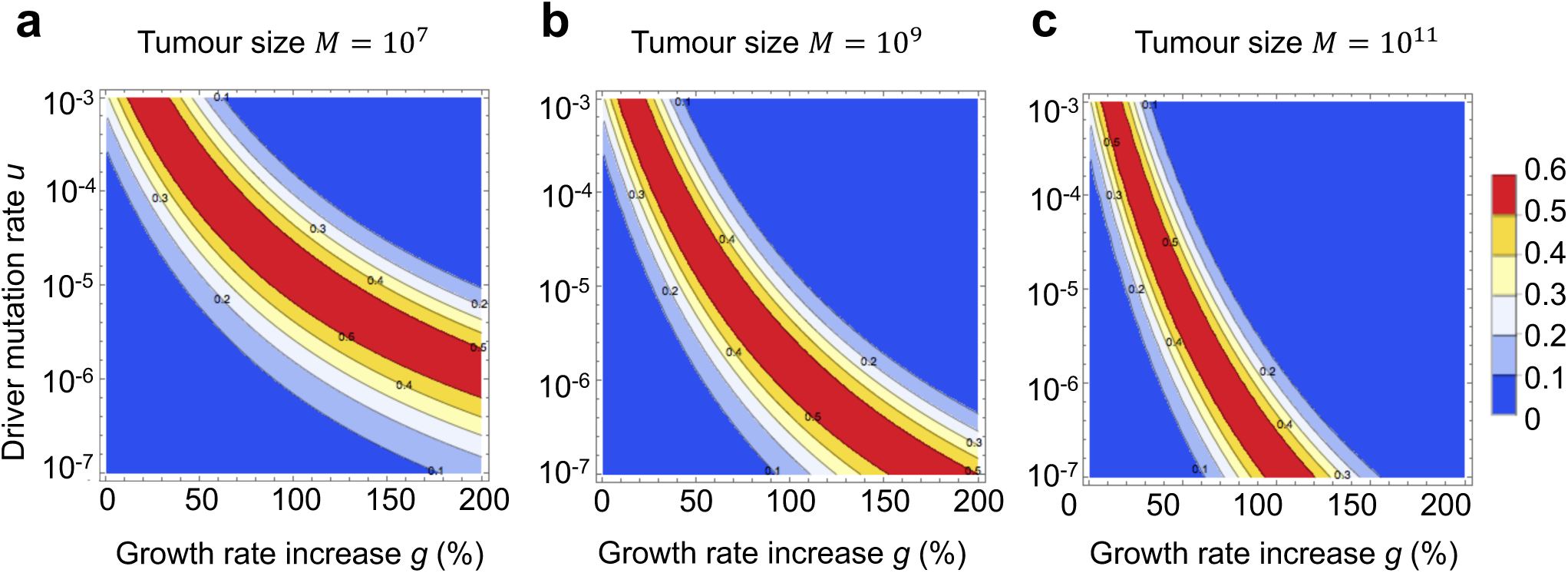
Probability of detection of a subclonal driver for a wide range of driver mutation rates and growth rate advantages is always below 60%. Contour plots depict the probability that a subclonal driver is in the detectable range (0.2 ≤ *f*_*sub*_ ≤ 0.8) and thus able to skew the distribution of mutation frequencies expected from neutral evolution, for **a,** small; **b,** intermediate; and **c,** large tumor size. Parameter values for moderately growing tumor *b*=*b*_1_=0.14, *r*=0.01. All rates are per day.

## DISCUSSION

In sum, the fact that no subclonal driver is present at intermediate frequencies cannot be taken as proof of neutral or “effectively” neutral evolution. It can simply be a consequence of population dynamics which creates only a short window during which the driver mutation can be detected but has not yet dominated the population.

Tarabichi et al.^19^ and McDonald and colleagues^20^ simulated tumor evolution in which they explicitly include selection, and showed that, even in models with selection, mutant allele frequency can exhibit the 1/*f* power law behavior, resulting in incorrect calling of neutrality. In response, Williams and colleagues^21^,^22^ argue that the example simulations from Tarabichi et al.^19^ and McDonald et al.^20^ that were incorrectly classified as neutral use extreme parameter values or correspond to either strong and early selection (a driver mutant quickly sweeps to fixation), or weak and late selection (driver mutants unable to reach detectable frequencies). In contrast, we show here that, for almost any driver mutation rate and selection strength, whenever we look at the mutant frequency spectrum of a tumor, it is likely either too early and the driver is present at a very low frequency, or it is already too late, and the driver is present in almost all cells of the tumor. Importantly, even if we manage to obtain the mutant frequency spectrum during the optimal window for detection, there is still significant chance (close to half) that the subclonal driver will not be in the detectable range.

Simulations of branching processes of cancer evolution for realistic tumor sizes and parameter values are computationally expensive. To circumvent that, studies often use small population sizes, death rate of cancer cells much smaller than the birth rate, and only examine a small set of different parameter values. In contrast, our mathematical results (formula 2) can be quickly evaluated for realistic parameter values, including all biologically plausible values of selection, mutation, birth and death rates, and population size. Furthermore, our results explain why the deviation of the mutant allele frequency from the 1/*f* power law in an intermediate frequency range is not a sensitive statistic for detecting subclonal selection in models of exponentially growing cancer populations: mutational frequency distribution of a subclonal driver is convex and thus always biased toward 0% and 100% frequency.

Our conclusion is relevant not only to cancer but more generally to the problem of distinguishing driver and passenger mutations when an exponentially-expanding subpopulation of fitter cells coexist with “wild-type” cells, such as exponentially growing bacterial populations acquiring *de novo* resistance to antibiotics or adapting to a new environment.

## MATERIALS AND METHODS

### Frequency distribution of a driver subclone

We study a two-type continuous time branching process that starts with a single type-0 cell. With rate *b*, type-0 cells divide into two identical daughter cells. Death rate of type-0 cells is *d*, with *b > d*. In addition, type-0 cells can receive an additional driver mutation with rate *u*. We will assume that the driver mutation rate is very small, on the order of *u ∼* 10^*-*5^ (per day). Cells with the additional driver divide with rate *b*_1_ and die with rate *d*_1_, again with *b*_1_ *> d*_1_. The net growth rate of cells with the additional driver, *r*_1_ = *b*_1_ *-d*_1_, is greater than the net growth rate of type-0 cells, *r* = *b-d*. We will denote the ratio of the two growth rates *r*_1_ and *r* by *c* = *r*_1_*/r >* 1. Let *X*_0_ be the number of type-0 cells at the appearance of the first successful cell with a driver (whose progeny survives stochastic fluctuations). The progeny of this cell forms the type-1 population. Total population size is the sum of type-0 and type-1 cells.

We are interested in the probability distribution of the fraction *f*_*sub*_ of type-1 cells in the population when total population size is *M*. Typical size of *M* that we will consider is 10^8^ *-*10^9^ cells. If we let *X* be the size of the type-0 population and *Y* the size of the type-1 population when total population size is *M*, then *f*_*sub*_ = *Y /M* and *M* = *X* + *Y*.

Survival probability of a cell with the additional driver mutation is *r*_1_*/b*_1_ = *cr/b*_1_. Thus the "successful" driver mutation rate (the rate at which driver cells with surviving progeny are produced) is *u*_*s*_ = (*cr/b*_1_)*u*. On the other hand, we have shown before that the arrival of mutations which appear with rate *u*_*s*_ in type-0 cells, can be viewed as a Poisson process with rate *u*_*s*_*/r* on the size of the type-0 population^24^. Thus the size of the type-0 population when the first type-1 cell appears, *X*_0_, is exponentially distributed with rate (*c/b*_1_)*u*. Thus *X*_0_ will be of the order of *b*_1_*/*(*cu*), which is typically much larger than 1, but much smaller than *M*.

We will measure time from the appearance of the first type-1 cell. Let *t* be the time when total population size is *M*. Since *X*_0_ is very large, the population of type-0 cells at time *t* can be well-approximated by *X ≈ e*^*rt*^*X*_0_. On the other hand, since type-1 cells started a surviving population at time 0 with a single cell, for the the population of type-1 cells we have^24^ *Y → V*_1_*e*^*crt*^ for large *t*, where *V*_1_ is an exponentially distributed random variable with rate *cr/b*_1_. In other words, *Y ≈ V*_1_(*X/X*_0_)^*c*^. It follows that

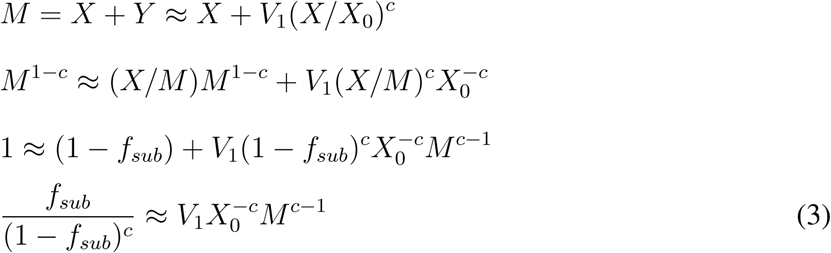

Here we used the fact that *X/M* =1 *- f*_*sub*_. On the other hand,

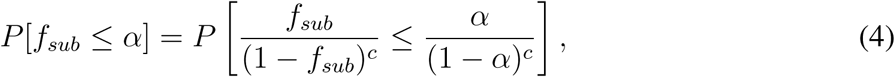

since *x/*(1 *- x*)^*c*^ is a function that increases as *x* increases from 0 to 1. Thus from (1) and (2) we have

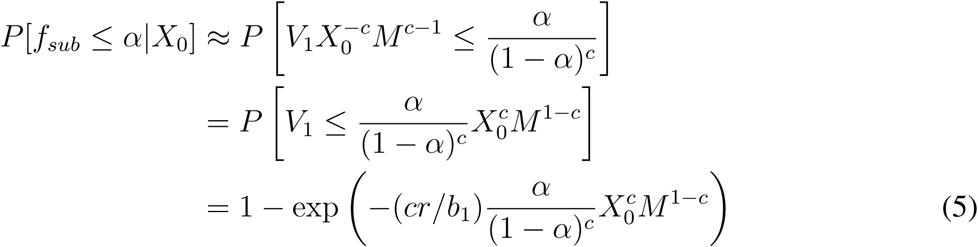

Finally we have

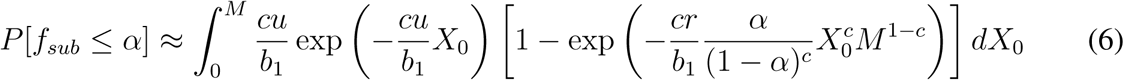

We show the excellent agreement of formula (6) and exact computer simulations of the process in Fig. 4.

**Figure 4.**
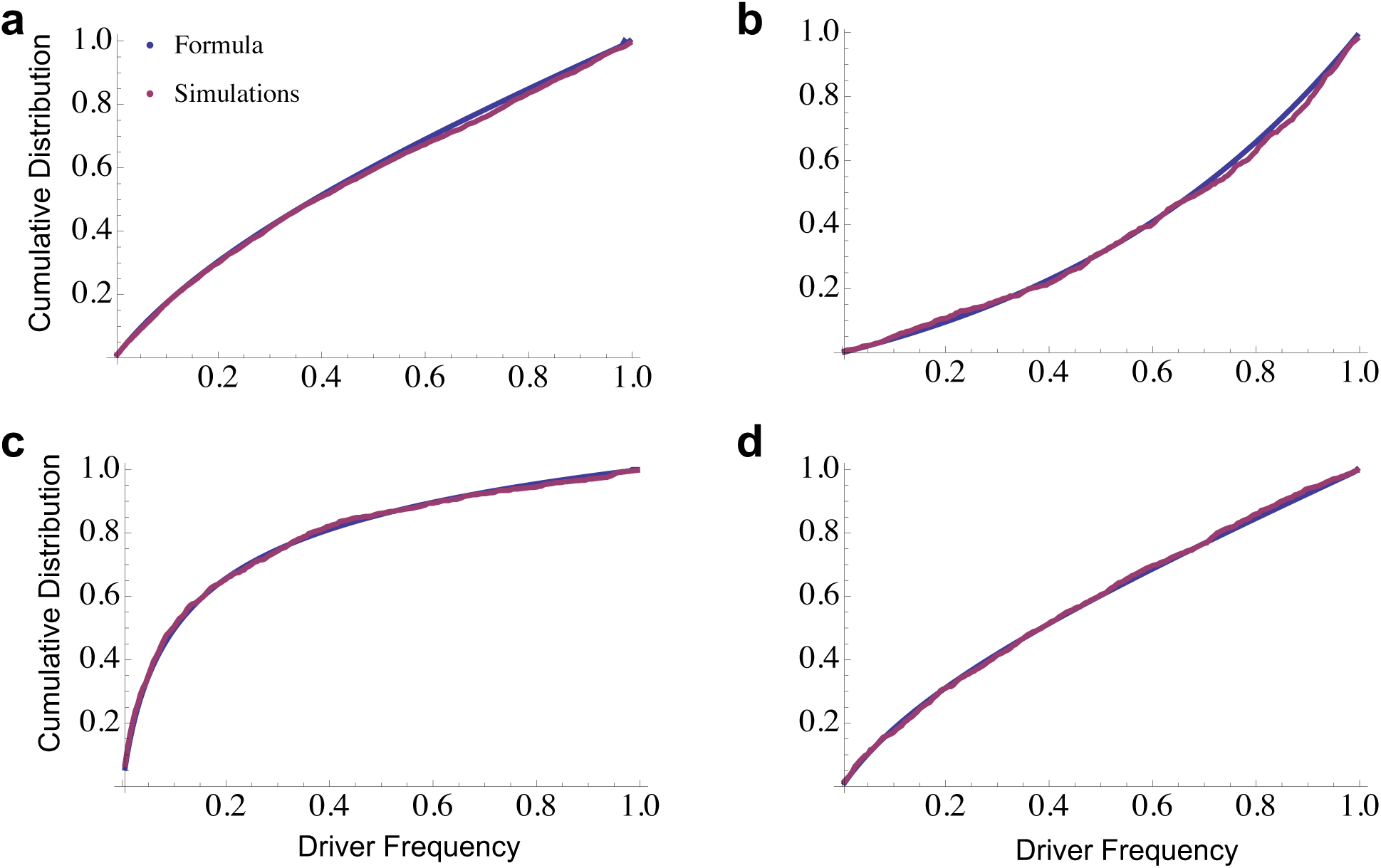
Comparison of formula for the cumulative distribution of driver frequency and exact computer simulations. On the y-axis we plot the probability that driver frequency is below a particular value. Parameters: **a,** *b*=*b*_1_=0.14, *d*=0.13, *c*=1.7, *u*=10^-5^, *M*=10^8^; **b,** *b*=*b*_1_=0.14, *d*=0.13, *c*=1.5, *u*=10^-4^, *M*=10^8^; **c,** *b*=*b*_1_=0.14, *d*=0.13, *c*=1.5, *u*=10^-5^, *M*=10^8^; **d,** *b*=*b*_1_=0.14, *d*=0.17, *c*=1.9, *u*=10^-5^, *M*=10^8^.

Let

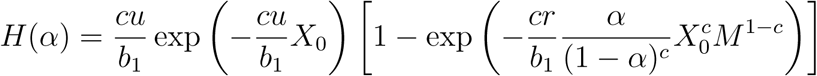

To calculate the probability density function, *f* (*?*) for the frequency of subclonal driver, we note that

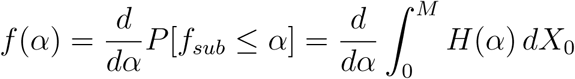

Using Leibniz’s rule we obtain

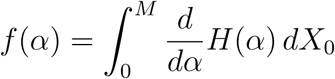

Finally, probability density function for the frequency of a subclonal driver is given by

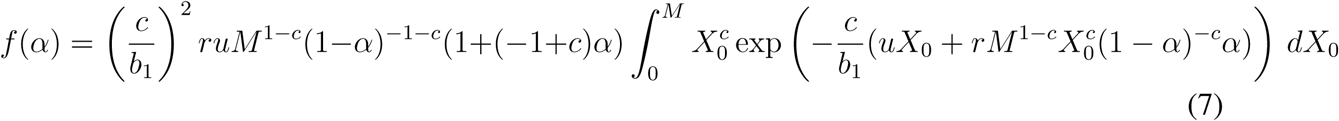

### Estimating driver mutation rate

Estimate for driver mutation rate *u ∼* 10^*-*5^ used in Fig. 2 comes from Bozic et al. ^18^ In that paper, it was estimated that there are 377 driver genes in the human genome, and an average of 90 positions per driver gene that, if mutated, will result in a functional driver mutation. In addition, it was assumed that the mutation rate per basepair per cell division was 5 *·* 10^*-*10^, leading to a driver mutation rate on the order of 10^*-*5^ per cell division.

Since then, new estimates have become available for both the number of driver genes and the point mutation rate in tumors. For example, Vogelstein et al. ^9^ use mutation patterns to estimate that there are 138 driver genes discovered so far. Similarly, Davoli et al.^10^ use analyzed patterns of mutational signatures in tumors and estimate 570 driver genes. Lawrence et al.^12^ use mutation frequencies and estimate 219 driver genes. They also perform a saturation analysis and show that many new candidate cancer genes remain to be discovered beyond those they report. Recently, Martincorena et al. ^11^ use dN/dS ratio to determine genes under positive selection in cancer and estimate 203 driver genes. Based on the sum of these data, we set the upper bound on the number of driver genes to be 600.

On the other hand, if we only focus on strong drivers in a single cancer type, such as colorectal, the number of genes is significantly smaller. For example, Martincorena et al. ^11^ report 28 genes under significant positive selection in colorectal cancer. Thus we will set the lower bound on the number of significant driver genes of a single cancer type to 20.

Blokzijl et al.^25^ estimate that *∼* 40 mutations accumulate per year in the genome of multiple human tissues, including the small intestine, colon and liver, leading to a mutation rate of 0.1/day per genome or *∼* 4 *·* 10^*-*11^ per basepair per day. This will be our lower bound for the point mutation rate. Recently, Werner and Sottoriva^26^ used the change in the mean and variance of the mutational burden with age in healthy human tissues and estimated mutation rate in the colon and small intestine to be *∼* 4 * 10^*-*9^ per basepair per cell division. Assuming the value they used for time between stem cell divisions of one week, this leads to a mutation rate of *∼* 6 *\** 10^*-*10^ per base-pair per day. Mutation rate in cancer can be increased 10-100 fold compared to normal tissues^27^, so we set the upper bound for point mutation rate to *∼* 100*\** 6*·*10^*-*10^ = 6*·*10^*-*8^ per base pair per day.

We obtain an upper bound for the driver mutation rate by multiplying the upper bounds for the number of driver genes and point mutation rate with the average number of driver positions, leading to *u*_*U*_ = 600 *\** 6 10^*-*8^ *\** 90 *∼* 10^*-*3^ per day.

Multiplying our lower bounds for the number of driver genes and point mutation rate with the average number of driver positions leads to the lower bound for the driver mutation rate *u*_*L*_ = 20 *\** 4 *·*10^*-*11^ *\** 90 *∼* 10^*-*7^.

## Subclonal driver frequency is biased towards 0 and 1 over a large range of parameter values

Using formula (6), we numerically evaluate *P* [0.2 *< f*_*sub*_ *≤*0.8] = *F* (0.8) *- F* (0.2) for the following ranges of parameters: ratio of net growth rates of cells with and without the driver, *c*, between 1.01 and 3 (i.e. relative growth rate advantage of a driver, *g*, between 1% and 200%); driver mutation rate, *u*, between 10^*-*7^ and 10^*-*3^ per day; final tumor size, *M*, between 10^7^ and 10^11^ cells; division rate of cells with the driver, *b*_1_, between 0.1 and 1 per day; and net growth rate of tumor cells, *r*, between 0.001 and 0.1 per day. These ranges are wide and include all meaningful parameter values.

We formulate a grid by taking 100 equally-spaced values for each parameter within its defined range (parameter values for driver mutation rate *u*, tumor size *M* and net growth rate of tumor cells *r* are equally spaced in log space). We exhaustively evaluate all points on this 5-dimensional grid (100^5^ = 10^10^ parameter value combinations). We find that *P* [0.2 *< f*_*sub*_ *≤*0.8] *<* 0.6 holds everywhere, and that the frequency of a subclonal driver is always biased toward 0 and 1.

